# Profiling embryonic stem cell differentiation by MALDI TOF mass spectrometry: development of a reproducible and robust sample preparation workflow

**DOI:** 10.1101/536664

**Authors:** Rachel E. Heap, Anna Segarra Fas, Alasdair P. Blain, Greg M. Findlay, Matthias Trost

## Abstract

MALDI TOF mass spectrometry (MS) is widely used to characterize and biotype bacterial samples, but a complimentary method for profiling of mammalian cells is still underdeveloped. Current approaches vary dramatically in their sample preparation methods and are not suitable for high-throughput studies. In this work, we present a universal workflow for mammalian cell MALDI TOF MS analysis and apply it to distinguish ground-state naïve and differentiating mouse embryonic stem cells (mESCs), which can be used as a model for drug discovery. We employed a systematic approach testing many parameters to evaluate how efficiently and reproducibly each method extracted unique mass features from four different human cell lines. This data enabled us to develop a unique mammalian cell MALDI TOF workflow involving a freeze-thaw cycle, methanol fixing and CHCA matrix to generate spectra that robustly phenotype different cell lines and are highly reproducible in peak identification across replicate spectra. We applied our optimised workflow to distinguish naïve and differentiating populations using multivariate analysis and reproducibly identifying unique features. We were also able to demonstrate the compatibility of our optimised method for current automated liquid handling technologies. Consequently, our MALDI TOF MS profiling method enables identification of unique features and robust phenotyping of mESC differentiation in under 1 hour from culture to analysis, which is significantly faster and cheaper when compared with conventional methods such as qPCR. This method has the potential to be automated and can in the future be applied to profile other cell types and expanded towards cellular MALDI TOF MS screening assays.

## Introduction

Matrix-assisted laser desorption/ionization time of flight mass spectrometry (MALDI TOF MS) is an versatile technique with many different applications ranging from protein identification by peptide mass fingerprinting and small molecule analysis to imaging of tissues.^1–3^ Although conventionally considered a low-throughput technology, recent advances in MS and liquid handling technologies and liquid handling tools has enabled MALDI TOF MS to emerge as a powerful tool for label-free high-throughput screening (HTS) within both the pharmaceutical industry and academic sphere.^4–6^ This platform is already well established for *in vitro* assays to monitor post-translational modifications such as ubiquitylation,^7,8^ phosphorylation^9,10^ and methylation,^11,12^ as the read-out is relatively simple with often just a single substrate and product. Similar to MALDI, laser desorption ionization can also be combined with self-assembled monolayers (SAMs), also known as SAMDI.^13,14^ Here, substrates are first immobilised on a surface before treatment with an enzyme, thus determining activity and kinetic parameters. Interestingly, SAMDI has been shown to be not only compatible with peptide substrates for protein specificity,^15,16^ but also carbohydrates and glycosyltransferase activity.^17^

Whole cell analysis or cellular assays for evaluating compound efficacy affecting a cellular phenotype presents an interesting challenge for MALDI TOF MS analysis as the system becomes inherently more complex. A well-established application for whole cell MALDI TOF MS is the profiling of micro-organisms, also known as biotyping.^18,19^ Profiling of protein biomarkers specific to a bacterial taxonomy by MALDI TOF MS was first performed by Claydon *et al* and enabled reproducible and robust identification of gram-positive and gram-negative species.^20^ Since then, bacterial genera have been identified through various approaches from spectral mass fingerprinting, to more complex approaches that involve comparing peaks identified in MALDI spectra to predictive masses from proteomic and genomic data sets.^21,22^ This in turn enabled the generation of reference protein databases that list biomarkers specific to different bacterial species.^23^ Combined with automated spectral acquisition and novel algorithms to tackle data analysis, biotyping has become a powerful, high-throughput tool for rapidly profiling bacterial genera in both academic and clinical settings.^24^ However, inter-lab studies revealed surprising discrepancies in *E. coli* fingerprints as experimental variables such as sample preparation and instrument parameters can affect spectral quality and reproducibility.^25,26^ Several studies have therefore scrutinised sample preparation methods for bacterial biotyping, looking at solvent extraction or direct analysis, sample handling and also matrix choice affects spectra quality with the aim of developing a standardised method to enable universal identification of micro-organisms.^27–29^

While bacterial biotyping has been very successful and has become a standard tool in the clinic, profiling of mammalian cells by MALDI TOF MS has not yet reached this level. High-resolution MALDI-FT-ICR mass spectrometry has been used by Sweedler *et al* to characterise lipids within 30,000 individual rodent cerebellar cells.^30^ This study enabled the identification of 520 lipid features and classification of neuron-like and astrocyte-like cells, thus allowing lipid profiles to be assigned to particular cellular functions. Characterising the protein signatures of mammalian cells by MALDI-MS is less common when compared with lipid analysis, however it has been used for phenotypic screening of human cancer cell lines,^31,32^ identification of cells within a co-culture^33,34^ or tissues^35^ and detection of transient changes within a specific cell type, such as immune cells.^36–39^ However, many of these studies list dramatically different experimental procedures with several being adapted from existing biotyping protocols. The huge range of experimental parameters could therefore be problematic for translation of published assays to the pharmaceutical industry.

To address the variation in experimental workflows we have systematically tested different methods at key steps in preparing mammalian cells for whole cell MALDI TOF MS analysis. We have generated a robust and sensitive sample preparation workflow by studying four commonly used human cell lines, followed by application of our final method to a pharmacologically controlled biological system, where we applied our optimised method to profile differences between naïve ground-state mouse embryonic stem cells (mESCs) and those undergoing differentiation. Thus, we have established a sample preparation method that is highly reproducible, robust and sensitive with respect to both biological and experimental variances and would be suitable for expansion to a HTS platform.

## Materials and Methods

### Human cell line culture

HEK293 and U2OS cell lines were cultured in DMEM media supplemented with 10% FBS, 1% pen/strep and 1% L-glutamine. MCF7 and THP-1 cells were cultured in RPMI-1640 media supplemented with the same and 50 µM β-mercaptoethanol was added to THP-1 cells. Adherent cell lines were lifted from 10 cm culture plates by addition of trypsin-EDTA solution. All cell lines were incubated in a controlled atmosphere at 5% CO_2_ and 37°C. Cells were harvested and centrifuged at 300 xg for 3 minutes before resuspension in PBS and counted using a haemocytometer. Cells were then aliquoted at a concentration 1 × 10^6^ into 1.5 mL microtubes and centrifuged at 300 xg, 4 °C for 10 minutes.

### Mouse Embryonic Stem Cell (mESC) Culture

CGR8 mESCs were cultured in 0.1% gelatin [w/v] coated plates in N2B27 medium (DMEM/F12-Neurobasal (1:1), 0.5% N2, 1% B27 (ThermoFisher Scientific), 1% L-glutamine, 100 μM β-mercaptoethanol) containing “2i”,^40^ 3 μM CHIR99021 (Axon Medchem) and 1 μM PD0325901, in a controlled atmosphere at 5% CO_2_ and 37°C. To induce multi-lineage differentiation,^41^ cells were plated at 4 × 10^4^ cells/cm^2^ in N2B27 medium without CHIR99021 and PD0325901 and incubated for 48h at 5% CO_2_ and 37°C.

### RNA extraction and qPCR

Total RNA extraction was performed by a column-based system (Omega) and then subjected to reverse transcription using iScript reverse transcriptase (Bio-Rad) according to the manufacturer’s guidelines. qPCR reactions were carried out using SYBR^®^ Premix Ex Taq™ II Supermix (Takara) in a CFX384 real-time PCR system (Bio-Rad). Samples were analysed for gene expression in 2i release conditions relative to 2i medium culture using the ΔΔCt method, and GAPDH expression was analysed as a loading control. Data from three independent biological replicates, with two technical replicates for each, were analysed in Excel software (Microsoft) and plotted in GraphPad Prism v.6.00 software (GraphPad). Primers used are listed in Table S-1. Statistical significance was determined using an unpaired Student’s t test, and significant differences were considered when p < 0.05.

### Cell microscopy and diameter analysis

The four cell lines were measured for number and cell diameter by light microscopy using an Evos XL Core Cell Imaging System (Invitrogen). Optimal cell numbers were calculated by a cell titration, whose values are reported in Table S-2, and these concentrations were used for subsequent experiments. To assess permeability, cell pellets were resuspended in PBS before mixing 1:1 with trypan blue. Trypan blue positive cells were then automatically counted using the same microscope to calculate cell viability. For mESC phenotype visualization, brightfield light microscopy was used in a Leica DM IL LED microscope at 10X magnification.

### BCA Protein quantitation

Cells were titrated from 300,000 to 9,000 in a 96 well plate format. BCA reagent (Pierce) was then prepared to the manufacturer’s instruction by missing Reagent A and Reagent B at a ratio of 50:1 respectively. 20 µL of the mixed reagent was added to 180 µL of sample and incubated at 37 °C for 30 minutes. The plate was then read on plate reader measuring absorbance at 562 nm and protein concentration calculated from these values.

### Sample preparation for MALDI TOF MS analysis

Cell pellets were processed in one of three ways:

(a) Direct analysis where cell pellets were washed twice with PBS and centrifuged at 300 xg, 4 °C for 10 minutes. Cell pellets were then resuspended in 0.1% TFA before subsequent spotting.

(b) Cell pellets were snap frozen on dry ice and stored at −80°C until required. Thawed cell pellets were centrifuged at 300 xg, 4 °C for 10 minutes before either being washed 1x with PBS or fixed in 4% paraformaldehyde solution or methanol on ice. Cell suspensions were then centrifuged at 300 xg, 4°C for 10 minutes before being resuspended in 0.1% TFA.

(c) Cell pellets were washed twice with PBS and centrifuged at 300 xg, 4 °C for 10 minutes. Cell pellets were then either washed 1x with PBS or fixed in 4% paraformaldehyde solution or methanol on ice. Cell suspensions were then centrifuged at 300 xg, 4°C for 10 minutes before being resuspended in 0.1% snap frozen on dry ice and stored at −80°C until required.

### Matrix preparation and spotting

Sinapinic Acid (SA), α-cyano-4-hydroxycinnamic acid (CHCA) and dihydroxybenzoic acid (DHB) were used as matrices for all MALDI TOF cellular analysis. All matrix solutions were prepared in 50% ACN, 0.1% TFA at varying concentrations and ratios of matrix solute: 2.5, 10, 20 mg/mL or saturated. For manual deposition cell suspensions were mixed at a 1:1 ratio with matrix solution and 1 µL spotted onto a ground steel MALDI target before ambient drying.

Automated target spotting was performed using a Mosquito liquid handling robot (TTP Labtech) by first mixing 1200 nL of matrix solution with 1200 nL of whole cell samples before subsequent deposition of 200 nL of the matrix: sample mixture an AnchorChip MALDI. The target allowed to ambient dry before analysis.

### MALDI TOF MS analysis

A RapifleX PharmaPulse MALDI TOF/TOF mass spectrometer (Bruker Daltonics) equipped with a Smartbeam 3D laser was used in positive ion mode with Compass 2.0 control for all data acquisition. Samples were acquired in automatic mode (AutoXecute; Bruker Daltonics), totalling 10,000 shots at a 10-kHz frequency per spot. Full MALDI-TOF MS parameters can be found in supplementary methods. MALDI TOF data processed by the FlexAnalysis 4.0 software where a peak picking threshold of 3 S/N was set before being exported as a .csv file using FlexAnalysis Batch Process (Compass 2.0) and further processed in Microsoft Excel and/or Perseus.^42^ Spectra based PCA plots were generated using ClinPro Tools (Bruker Daltonics). Data was then formatted using both GraphPad Prism 7.0 and Adobe Illustrator.

### Bootstrap statistical analysis

Hierarchical cluster analysis was performed on all biological and technical replicates of the two conditions using the R package pvclust, with multiscale bootstrap resampling of 10,000 iterations used to assess statistical significance by approximately unbiased p-values.^43,44^ Clustering was implemented using the average agglomeration method with a correlation distance metric. The results are presented alongside a z-score heatmap of detected mass features.^45^

## Results and Discussion

### Optimizing the workflow for mammalian cell MALDI TOF MS

In order to optimize the sample preparation for whole cell MALDI TOF MS, we focused initially on four different human cell lines (U2OS, MCF7, THP1 and HEK293) (Figure 1A). Cells were washed once with PBS, which was sufficient to remove the culture medium contaminants as high levels of foetal bovine serum (FBS) and salts from the culture medium affect MALDI TOF MS ionization.^36^ To determine optimal cell concentration, we spotted 25 to 20,000 cells on target. Surprisingly, there was a narrow window where good spectra could be acquired, with large numbers of cells on-target proving to be detrimental to ionization.

**Figure 1:**
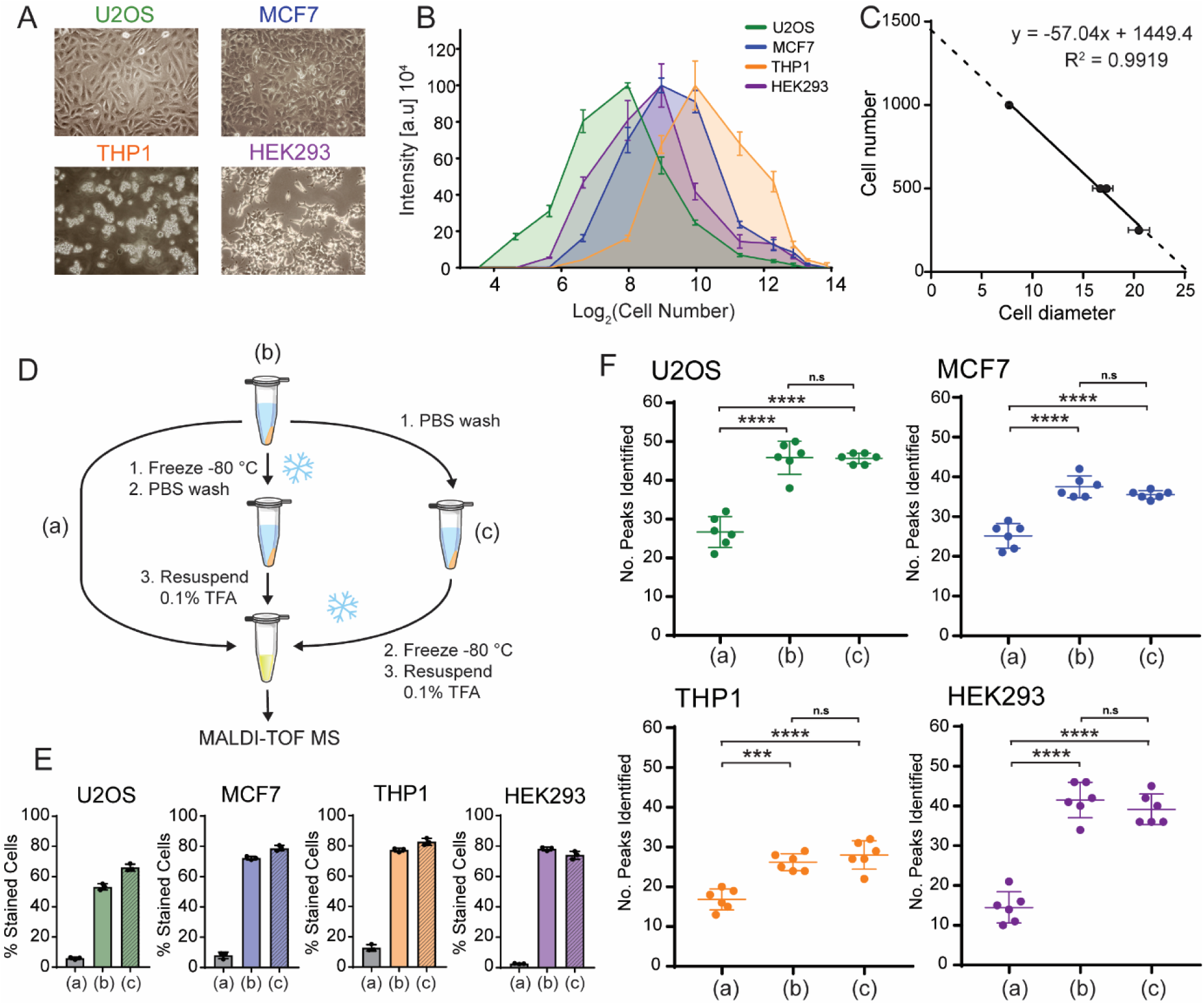
Optimizing cell numbers for whole cell MALDI TOF MS. (A) Light microscopy images of four human cell lines U2OS, MCF7, THP1 and HEK293. (B) 2D plot of normalized mass spectrum intensity at different cell numbers on target for each of the four cell lines. Plots have maxima indicating optimal cell numbers. (C) Plotting optimal cell numbers derived from (B) against their respective measured cell diameter shows a high correlation. Plots in (B) derived from 6 technical replicates. Error bars represent standard deviation. (D) Schematic showing workflow. (E) Trypan blue staining of the four cell lines shows that the freeze/thaw cycle significantly increases percentage of trypan blue positive cells and therefore cell membrane permeability. (F) Number of peaks identified across six technical replicates for each of the experimental workflows shown in (D). Data show that the number of features identified are significantly increased upon inclusion of a freeze/thaw cycle. Error bars represent standard deviation of six replicates. *** and **** represent p<0.001 and p<0.0001, respectively, student’s t-test.

Further to this, we observed that the best spectral intensity varied for each cell line (Figure 1B) and hypothesised that the number of ionisable biomolecules from cells was dependent on the cell size. This relationship was also consistent between protein concentration and cell number (Figure S1). Therefore, we measured the diameters of all four cells lines in solution (Table S-2) and plotted these values against the optimal cell number derived from the titration to identify an optimal cells number on-target for MALDI TOF MS analysis (Figure 1C). This generated a linear relationship with a very good correlation of R^2^ = 0.99 indicating that to obtain optimal and reproducible spectra from mammalian cells by MALDI TOF MS, cell numbers on target need to be optimised and this number is dependent on cell size. This generated a linear relationship with a very good correlation of R^2^ = 0.99 indicating that to obtain optimal and reproducible spectra from mammalian cells by MALDI TOF MS, cell numbers on target need to be optimised and this number is dependent on cell size.

We tested next if the biomolecules detected from mammalian cells derive from intact cells or if mild breakage of cells enhanced the occurrence of unique mass features. In our experience, harsh lysing conditions resulted in spectra that were less distinguishable, which has also been observed by lysing with increasing acidity.^37^ It was indicated before that freeze-thawing of cell pellets prior to MALDI TOF MS analysis may have beneficial effects with respect to number of features identified and overall spectral intensity.^33,46^ This is likely due to the freeze/thaw cycle permeating the cell membrane, thus allowing the cytoplasmic contents of the cells to become exposed and more easily ionised. We therefore decided to test whether a freeze/thaw cycle improved MALDI TOF MS analysis of mammalian cell lines and whether freezing before or after a wash with PBS affected sensitivity and spectral quality compared with direct analysis (Figure 1D).

Both methods of freeze/thawing permeated the cell membrane of about 50-80% of the cells (Figure 1E). This led to a significant increase in the number of peaks identified compared to “intact” cell samples (Figure 1F). As well as this, software analysis did not result in a significant difference between cells frozen before or after further treatment and manual inspection of spectra resulted in the same conclusion (Figure S2). We conclude that a freeze/thaw cycle is critical to improve MALDI TOF MS quality of mammalian cells as it increases the number of features identified. However, the order in which this step is performed, does not affect the final readout.

### Suitability of mammalian cell fixing techniques for whole cell MALDI TOF-MS

Next, we took inspiration from cell and tissue fixing techniques and examined how different fixing techniques influence the preparation of mammalian cells for MALDI TOF MS analysis. We hypothesised that the use of these techniques would enable preservation of a specific cellular phenotype and could be incorporated into a whole-cell MALDI TOF-MS compatible workflow. We chose to initially test formaldehyde and alcohol fixing methods, methanol specifically, as they have been used previously in MALDI imaging workflows.^47^ Initial experiments revealed that samples treated with 4% PFA generated spectra that were 5 – 10x less intense than other methods, (Figure S3-6) therefore we chose to continue by studying methanol fixation versus the previously reported PBS washes. We systematically evaluated how both these methods performed with respect to the number of identified peaks, quality of the acquired spectra, as well as technical reproducibility when analysed by MALDI TOF MS.

Both methanol and PBS washing steps distinguished each of the four cell lines by both manual spectra interrogation and principle component analysis (PCA) (Figure 2A-F) and, each extraction method was able to generate a unique set of peaks for each of the four cell lines, thus allowing classification of the different populations. As phenotype distinguishing peaks are often not the most intense features and peaks identified with a lower S/N and intensity are less likely to be quantified accurately, we therefore looked at how the relative intensity of peaks was distributed for the top 10 most intense peaks for each cell line (Figure 2G & H). This is important for high-throughput analysis, and both PBS and methanol treatment showed a generally even intensity distribution. Finally, and arguably most importantly, we tested at how reproducible peaks were identified over six technical replicates (Figures 2 I & J, S2-5). Methanol was the most consistent, with the majority of all peaks being identified in all six spectra, whereas PBS and water (pH 7) were slightly more variable. Taken together, our data suggests that methanol fixation is comparable if not superior with PBS washing for whole cell analysis and may be ideal for classification of subtle phenotypes.

**Figure 2:**
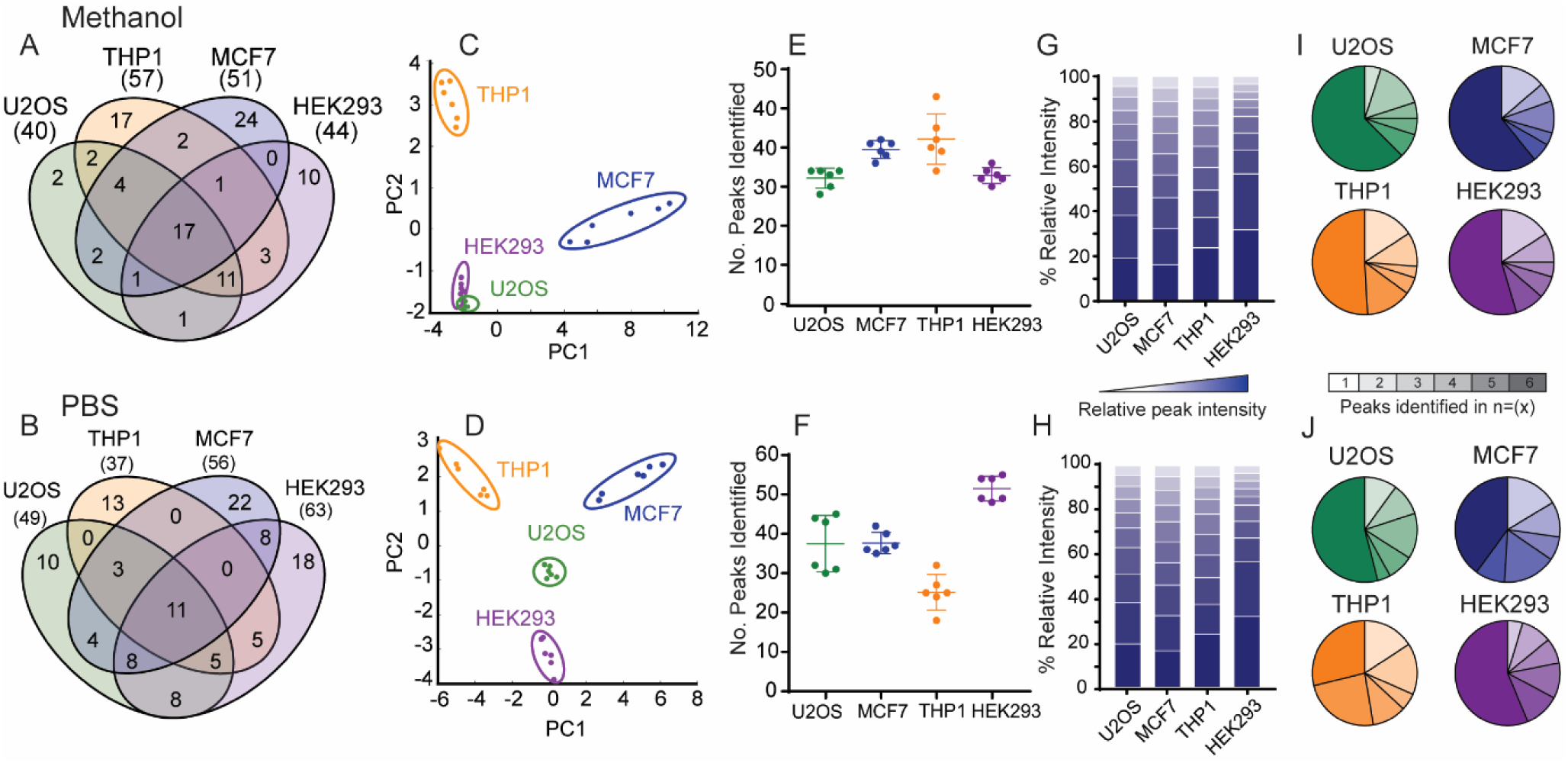
Methanol fixation and PBS washing are suitable for identifying reproducible features in whole cell MALDI TOF MS. (A & B) Number of identified unique features for any of the four cells after methanol fixation (A) and PBS washes (B). (C & D) PCA plots showing how multivariate analysis can distinguish each of the four cell lines after methanol fixation (C) and PBS washes (D). (E & F) Number of peaks identified for each cell line over six technical replicates. (G & H) Relative intensity distribution of the top 10 peaks identified over six technical replicates for each cell line. (I & J) Pie charts displaying the % of peaks identified within each of the six replicates per cell line.

### Choosing a suitable matrix for mammalian cell MALDI TOF MS

Next, we tested which type of matrix allows for the best MALDI TOF MS analysis of mammalian cells. The three matrices mostly used in MALDI TOF MS of proteins and peptides are sinapinic acid (SA), α-cyano-4-hydroxycinnamic acid (CHCA) and dihydroxybenzoic acid (DHB), which are often categorised for the analysis of proteins, peptides, and glycans, lipids and peptides, respectively. However, when analysing whole mammalian cells by MALDI TOF MS the origin of the biomolecules being ionised is often unknown, and we therefore hypothesised that the choice of matrix will have a significant influence on the resulting mass spectrum.

As expected, when each cell line sample was prepared with either saturated SA, CHCA or DHB, significantly different mass profiles of the same cell line were observed (Figure S7). Using DHB matrix resulted in more variable spectra over technical replicates, with the PCA analysis using ClinPro Tools software showing wider distances and grouping only three of the six technical replicate spectra for MCF7 cells (Figure S8A). Moreover, we could classify unique peaks to each of the matrices (Figure S8B), which indicates that different biomolecules are being ionised and mammalian cell profiles are matrix-dependent.

We then chose to look at how each matrix performs with respect to concentration and increasing laser power. In all four cell lines, CHCA performed significantly better with respect to both parameters. Although each matrix performed optimally at a saturated concentration, CHCA was able to yield much more informative spectra and with more peaks identified at a third of the concentration of DHB and SA. Following this, we observed that CHCA matrix was also able to ionise cellular biomolecules at much a lower laser power than DHB and SA. We were able to fit non-linear curves to these data, thereby identifying optimal minimal laser power and approximate saturation points of each matrix (Figure 3, Table 1). As CHCA is known for ionising peptides and small proteins, these data indicate that the biomolecules are either peptidic or small molecular weight proteins.

**Table 1.** Laser power parameters determined from non-linear regression curves shown in Figure 4B.

| Laser power parameters | U2OS |  |  | MCF7 |  |  | THP1 |  |  | HEK293 |  |  |
| --- | --- | --- | --- | --- | --- | --- | --- | --- | --- | --- | --- | --- |
| Matrix | CHCA | DHB | SA | CHCA | DHB | SA | CHCA | DHB | SA | CHCA | DHB | SA |
| Optimal laser power (%) | 38 | 83 | 70 | 32 | 79 | 73 | 35 | 71 | 76 | 31 | 73 | 78 |
| Saturation point (%) | 60 | ~90 | 90 | 60 | ~90 | 80 | 50 | ~90 | 80 | 60 | ~90 | 90 |

**Figure 3:**
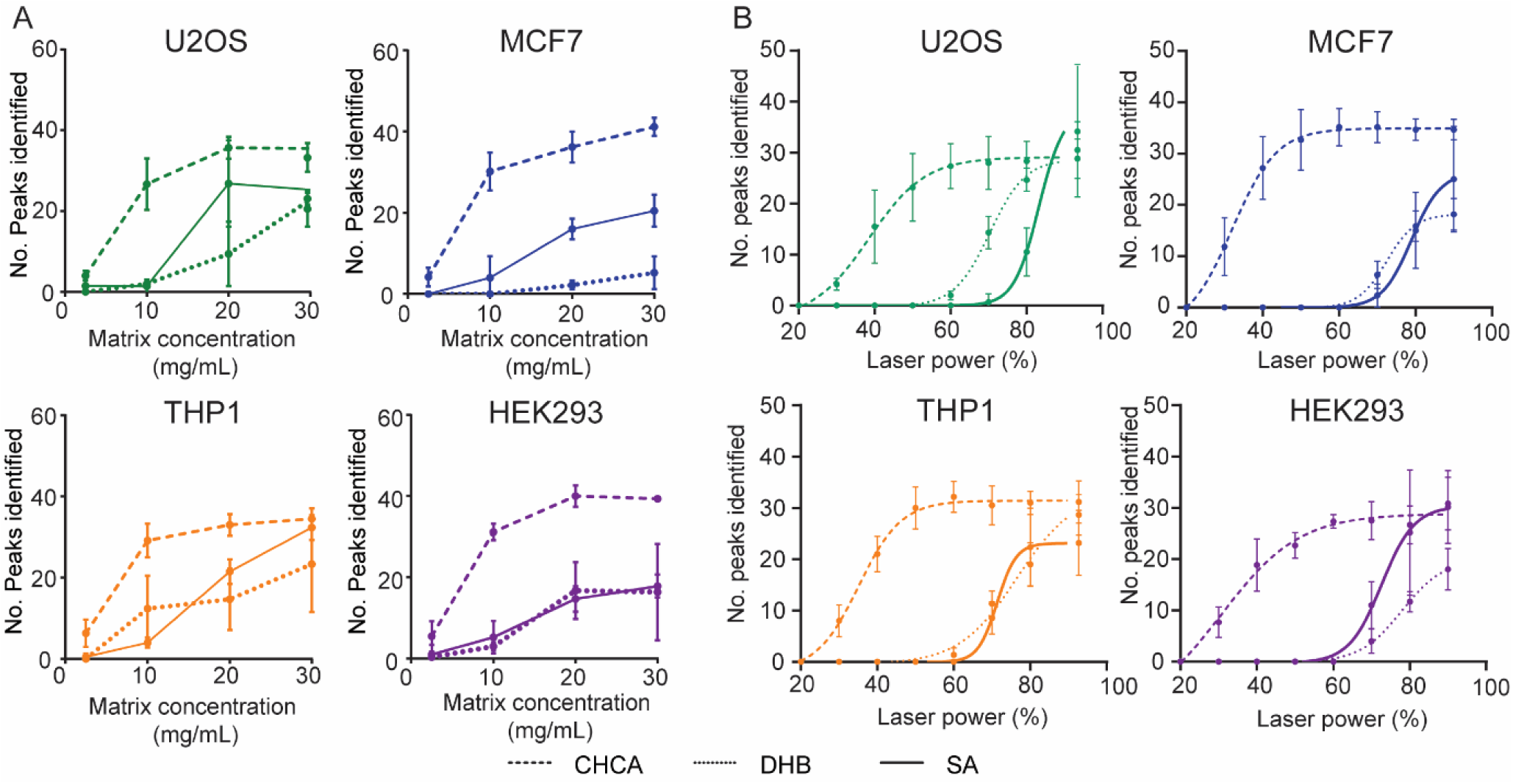
Evaluating matrix performance when analysing whole cell samples by MALDI TOF MS. (A) Dependence of matrix concentration from 2.5 – 30 mg/mL (saturated) on number of peaks identified for each of the four human cell lines and three matrices. (B) Laser power dependence of each of the three matrices for each of the four cell lines. Non-linear regressions were fitted with a Sigmoidal, 4PL curve with good fits of R^2^ >0.85 – 0.99. All error bars are given as standard deviations over six technical replicates.

We chose to take these data further to understand the dynamics of matrix behaviour in a mock screen and evaluate their performance across a 1536 target. We chose to use methanol treated THP-1 cells that were then mixed with each matrix using a Mosquito HTS and spotted in 200 nL aliquots. Each target was then analysed at the approximate saturation points described in Table 1: 60%, 80% and 90% for CHCA, SA and DHB, respectively. When compared a laser power of 60%, this corresponded to a laser energy fold change of 1.74x and 2.30x for 80% (DHB) and 90% (SA), respectively. From manual inspection as well as a positive MALDI response we determined a spotting accuracy of >96% for each target (Figure 4A). This infers that methanol-fixed whole-cell samples are compatible with current liquid handling technologies, as well as MALDI TOF-MS.

**Figure 4:**
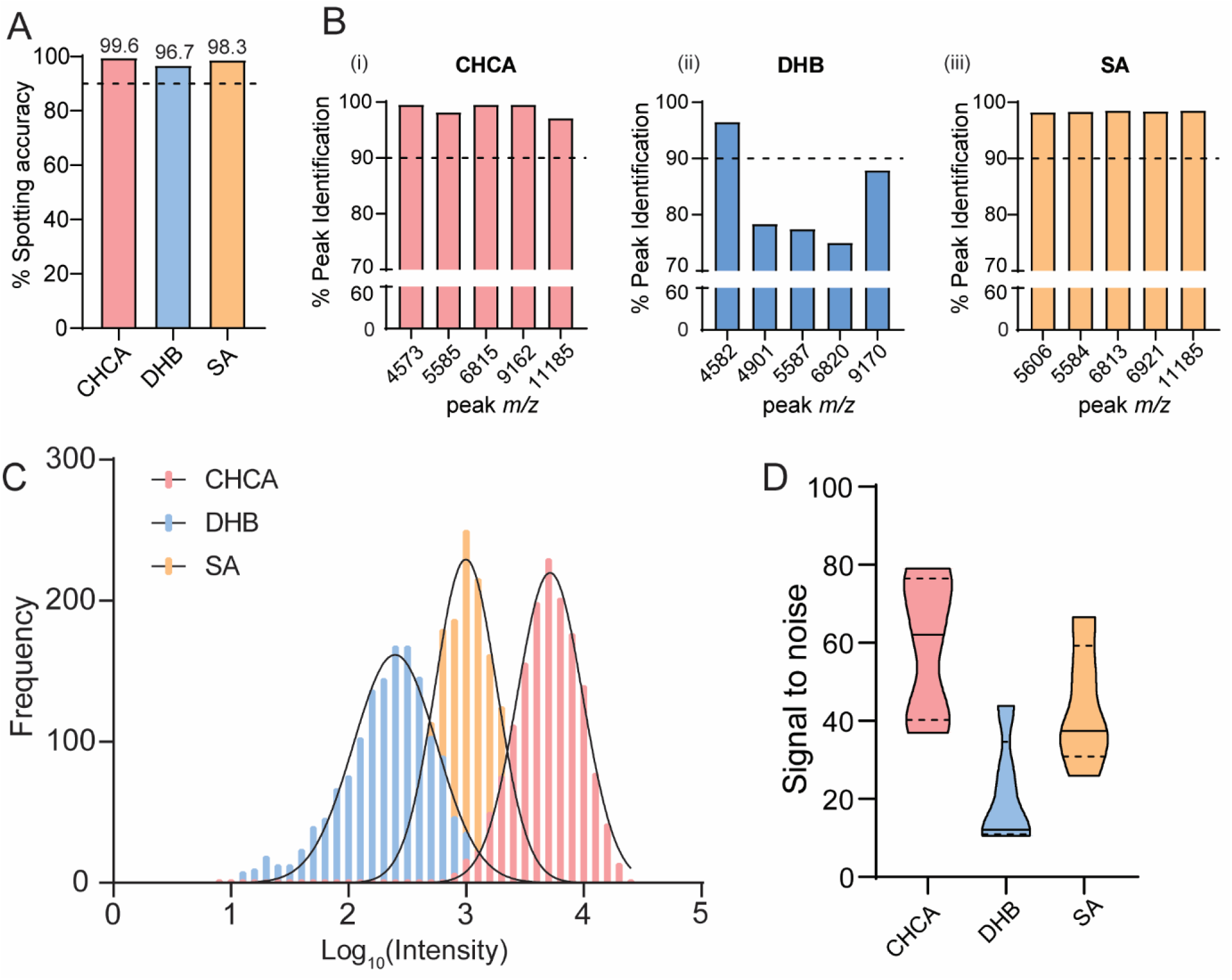
CHCA is the most suitable matrix for studies with a large sample number. (A) Mosquito spotting accuracy determined by positive signal spectra and manual inspection for each of the matrices. Percentage of 1536 spots. (B) Percentage peak identification of the top 5 most intense peaks over a 1536 target for each of the three matrices. (C) Frequency distribution of the top 5 most intense peak intensities of a 1536 target for each of the three matrices. (D) Violin plots showing distribution of the signal-to-noise of the top 5 most intense peaks over a 1536 target for each of the three matrices. Solid lines indicate median values and dotted lines the upper and lower quartiles.

We were also able to show that for CHCA and SA, the top five most intense features were identified in >98% of spots, showing robustness for high-throughput screening, whereas for samples spotted with DHB peak identification is much more variable (Figure 4B). This is somewhat expected as DHB crystallises into long, needle-like structures that produce a heterogeneous surface. This combined with an inherently heterogeneous sample such as fixed cells may account for this variability. Overall spectral intensity based on the top five peaks as well as signal to noise also varied significantly between the matrix conditions. Samples spotted with CHCA exhibited much greater spectral intensity compared to SA, and an almost 10-fold increase when compared with DHB (Figure 4C), as well as significantly better S/N for these top 5 features (Figure 4D). Interestingly, we observed different lens contamination patterns for each of the three matrices after accumulation of 3072 individual spectra with each matrix (Figure S9). Both DHB and SA yielded significant deposition of matrix onto the lens compared with CHCA. From our data set this does not appear to negatively impact whole cell ionisation, however we do suspect that prolonged ionisation and exposure to samples co-crystallised with either SA or DHB will impact studies that have a greater number of samples such as those in high throughput screens. From this data, we conclude that CHCA would be the matrix choice for whole cell analysis at a small and large scale due to its superiority across the parameters discussed above. However, we do report that for masses greater than >10,000 Da, peak resolution is significantly improved by using SA (Figure S10) and therefore may have a role to play in studies that identify significant features in this mass range. Together, these results show how phenotyping cells by MALDI-TOF MS can be scaled up to a high-throughput platform and still enable robust identification of cell specific features.

### MALDI TOF MS profiling of pharmacologically controlled stem cell differentiation

Finally, we wanted to apply our optimised workflow to phenotypically profile cells in an physiologically relevant system that is employed in drug screening and toxicity testing and that has been used as a drug discovery model.^48^ We used mouse embryonic stem cells (mESC) maintained in a naïve ground state pluripotency using the 2i kinase inhibitor system (PD0325901 and CHIR99021), which inhibit the kinases ERK1/2 and GSK3, respectively (Figure S11).^40,49^

Efficient exit from naïve ground state pluripotency towards differentiation upon 2i release was confirmed by suppressed mRNA expression of the naïve pluripotency factors *Nanog* and *Klf4*, and induction of the lineage priming/differentiation marker *Fgf5* (Figure 5A). As expected, the pluripotency factor Pou5f1/Oct4, which is expressed in both naïve and lineage primed mESC states, is not significantly altered upon acute 2i release (Figure 5A). Using MALDI TOF MS, we could robustly identify unique features to each population, as well as quantify changes in common peaks. For all spectra, the base peak was identified at *m/z* 4875, which made subsequent analysis simpler, as the raw spectral intensity can vary significantly from spot-to-spot (Figure 5D & Figure S12). Utilizing *m/z* 4875, as a normalizing control, we identified a number of peaks that were unique to 2i and 2i release, such as *m/z* 5566, 2i and 2i release conditions could be well differentiated as two populations by PCA (Figure 5B) and we observed good grouping of biological replicates. A similar distribution was identified when using a jackknife method (Figure S14, Table S3).^50^ To further understand how relative intensity of specific peaks changed across the three biological and five technical replicates, we generated a Z-score averaged based heatmap of detected mass features relative intensity (Figure 5C). For robustness, data were first filtered to include features that were identified in 30% of all technical replicates. A complete heatmap of all features is presented in Figure S15. This hierarchical clustering approach allowed us to look at the unique and common features combined across all independent biological and technical replicates. The three biological replicates clustered well together and 2i and 2i release conditions were separated efficiently and two discrete row clusters emerged – peaks that were up-regulated and those that were down-regulated upon release from PD0325901 and CHIR99021 inhibitors. This clustering was significant in the hierarchical dendogram (Figure S16) showing classification of the two different cell populations. Through this, we identified several peaks that changed significantly between the conditions and these can now be used as features of phenotypic screening of mESC differentiation (Table 2). Our MS approach has significant advantages over the conventional qPCR approach with respect to time, cost and automation possibilities. Using MALDI-TOF MS and liquid handling robots, samples could be processed in a 384 well plate format and analysed within one hour, approximately three times faster than using qPCR. Furthermore, only 1000 cells are required to phenotype their differentiation state, comparable with qPCR, but the consumable cost per sample is significantly reduced. Consumables for our MALDI-TOF MS assay are about £0.05-0.10 per sample, requiring only basic plastic ware, low solvent volume, CHCA matrix and stainless steel MALDI targets. These are significantly cheaper per sample compared to qPCR plate kits that typically cost >£2-5 per sample. Finally, our MALDI-TOF MS approach has the capability to be automated to a high-throughput scale using already established technologies such as the Mosquito HTS.

**Table 2.** Significant features identified in both 2i and 2i release spectra that exhibit a greater than 2 fold change.

| m/z | Relative fold increase |  |
| --- | --- | --- |
|  | 2i | 2i release |
| 2437 | x | 2.03 |
| 5438 | x | 2.70 |
| 7148 | x | 2.08 |
| 7412 | x | 8.82 |
| 3097 | 4.57 | x |
| 4628 | 2.34 | x |
| 6197 | 3.18 | x |
| 11174 | 2.86 | x |

**Figure 5:**
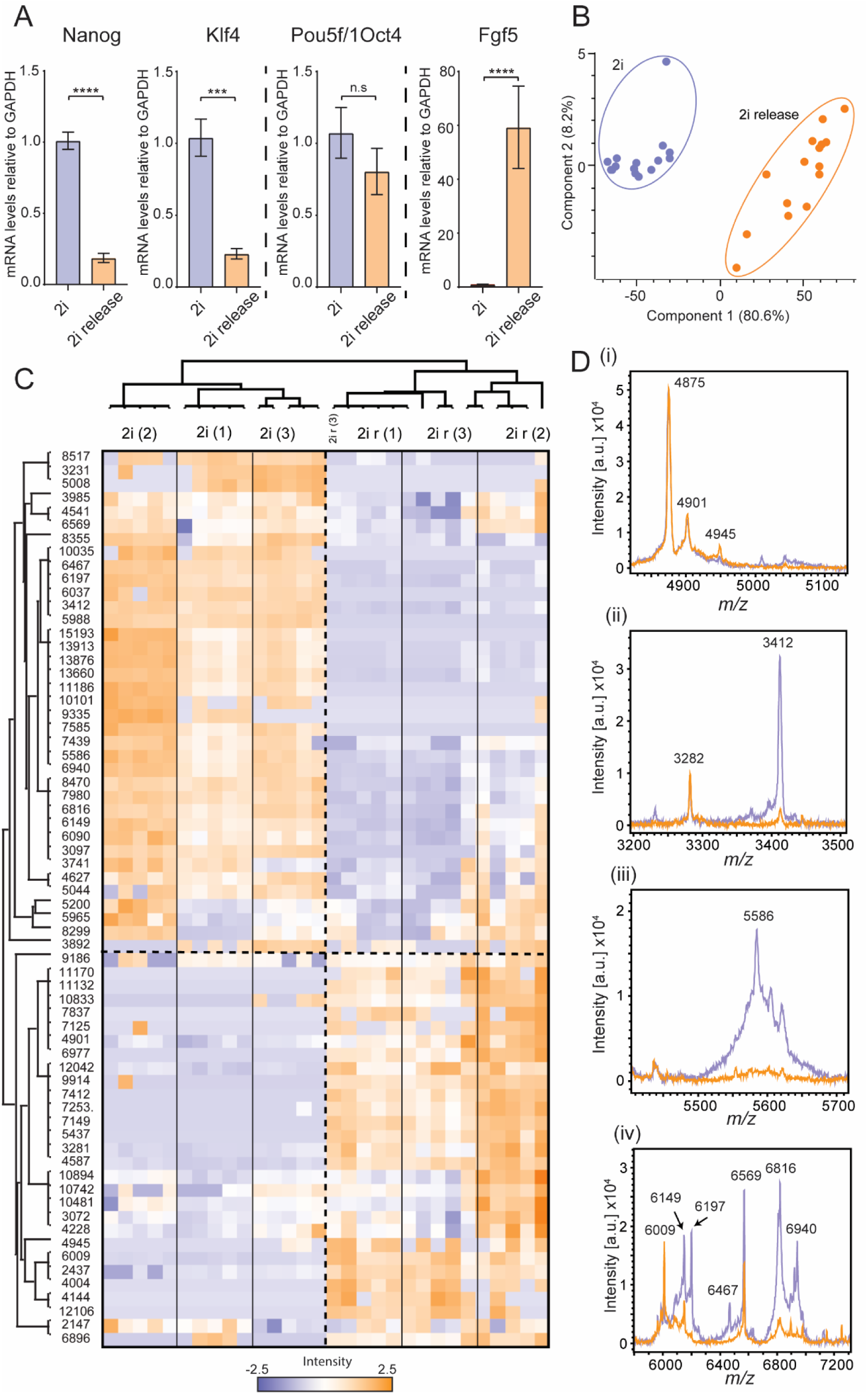
Whole cell MALDI TOF MS distinguishes between naïve and differentiating mESCs. (A) qPCR data showing changes in key mESC pluripotency and differentiation genes. Error bars represent standard deviation of two technical replicates from three biological replicates. *** and **** represent p<0.001 and p<0.0001, respectively, student’s t-test. (B) PCA plot showing how the two cell populations can be distinguished by multivariate analysis over three biological and five technical replicates. (C) Z-score heat map showing how each normalized peak intensity changes with respect to 2i and 2i release condition over three biological and five technical replicates. (D) Selected examples of mass spectral regions that showed changes between mESCs cultured in 2i and 2i release conditions where the base peak (4875 *m/z*) is of identical intensity, thus allowing relative intensity comparison.

## Conclusion

Due to its speed and its relative simplicity, MALDI TOF MS has become increasingly popular for the application of bacterial biotyping. However, a complimentary methodical approach to phenotypic screening of mammalian cells has not been well characterised. Here we presented a systematic study that explores initial sample handling, matrix choice and suitability of fixing techniques with whole cell MALDI TOF-MS analysis. We found that at all three steps had a profound impact on the resulting mass spectra and subsequent data analysis. We also applied a unique way of analysing the efficacy of each method by looking at not only spectral quality and observable changes but evaluating performance over technical replicate spots. This enabled us to gain a deeper understanding of how each step of the sample preparation impacts subsequent analysis and consequently an insight as to how each method would perform with higher throughput analyses. Our optimised method was validated by our observation of distinct MALDI TOF MS profiles for naïve ground state mESCs compared to differentiating mESCs in a pharmacologically controlled system. Using hierarchical clustering, we could visualize and identify a subset of peaks that are unique to each condition. We therefore present here a novel sample preparation method that enables robust, reproducible and rapid profiling of mammalian cells and is suitable for expansion to high-throughput platform.

## ASSOCIATED CONTENT

Supporting Information

Supplementary figures, tables and texts (PDF)

Complete Table of 2i and 2i release relative intensities (.xlsx)

## Author Contributions

The manuscript was written through contributions of all authors. / All authors have given approval to the final version of the manuscript.

## Notes

The authors declared no potential conflicts of interest with respect to the research, authorship, and/or publication of this article.

## ACKNOWLEDGMENT

We thank the Bruker Daltonics, particularly Meike Hamester, Anja Resemann and Astrid Erdmann, for their support. We thank Anetta Härtlova, Adam Moore, Shin Lim and Julien Peltier for help. The authors disclosed receipt of the following financial support for the research, authorship, and/or publication of this article: This work was funded by generous start-up funds from Newcastle University to MT, an iCASE studentship to REH by the BBSRC and Bruker Daltonics. GMF is funded by a Wellcome Trust/Royal Society Sir Henry Dale Fellowship, ASF is supported by a studentship from the MRC.

