## Supplementary tables and figures for "Profiling embryonic stem cell differentiation by MALDI TOF mass spectrometry: development of a reproducible and robust sample preparation workflow" (from another version of this preprint: version 1, 2019-02-04)

Supplementary Information

**Table S-1:** Statistical significance was determined using an unpaired Student's t test, and significant differences were considered when $p < 0.05$.

| Gene | Forward (5' to 3') | Reverse (5' to 3') |
|---|---|---|
| *Nanog* | CTCATCAATGCCTGCAGTTTTTCA | CTCCTCAGGGCCCTTGTCAGC |
| *Klf4* | ACACTTGTGACTATGCAGGCTGTG | TCCCAGTCACAGTGGTAAGGTTTC |
| *Oct4* | AGCTGCTGAAGCAGAAGAGG | AGATGGTGGTCTGGCTGAAC |
| *Fgf5* | GCTGTGTCTCAGGGGATTGT | CACTCTCGGCCTGTCTTTTC |
| *Dnmt3b* | CTGGCACCCTCTTCTTCATT | ATCCATAGTGCCTTGGGACC |
| *Gapdh* | CTCGTCCCGTAGACAAAA | TGAATTTGCCGTGAGTGG |

**Figure S-1: Freeze/thaw method spectra.** MALDI-TOF spectra of each of the four cell lines (U2OS, MCF7, THP1, HEK293) for the freeze/thaw methods: direct analysis (a), freeze/thaw first (b), freeze thaw after wash (c). Freeze/thaw cycle significantly increases number of features identified for each of the cell lines. The point at which this step is performed does not negatively affect spectral sensitivity or information.

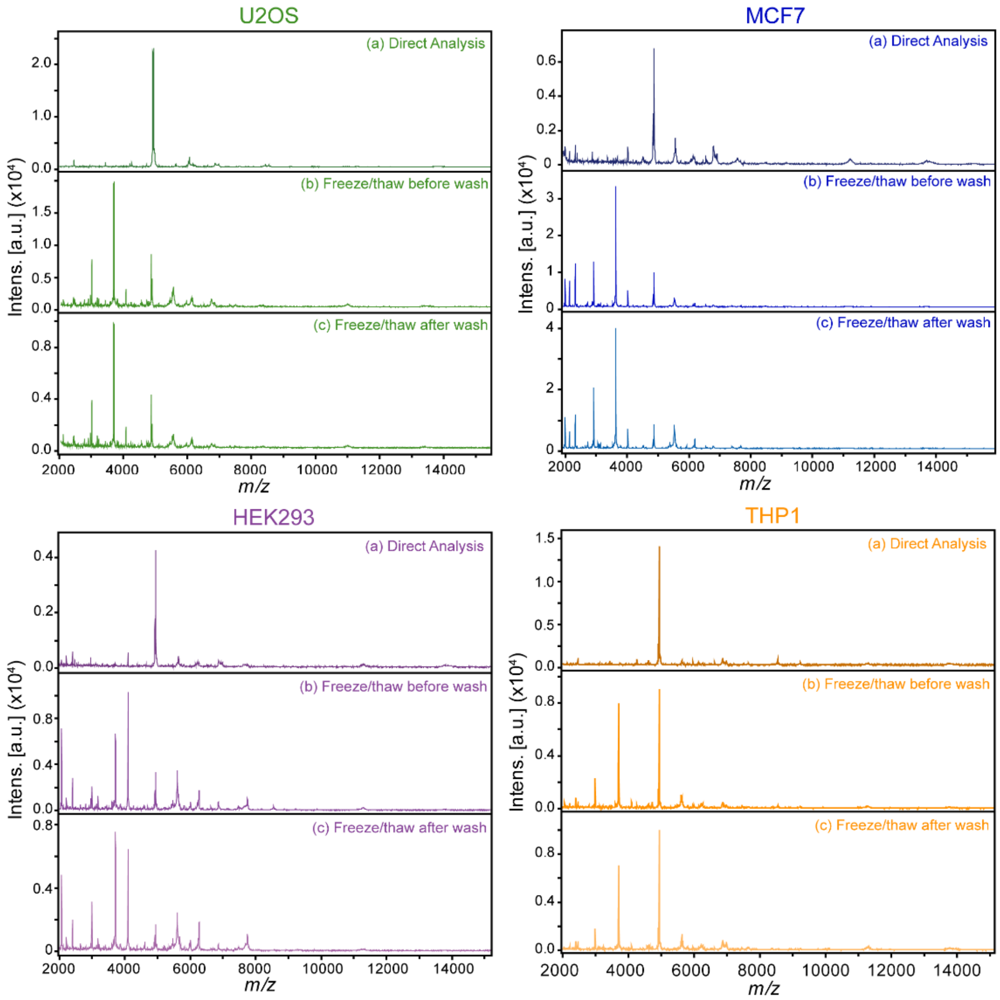

**Figure S-2: PBS wash spectra**. MALDI-TOF spectra of each of the four cell lines (U2OS, MCF7, THP1, HEK293) that were washed with PBS before subsequent analysis. For each cell line good spectra were acquired with unique profiles being generated for each.

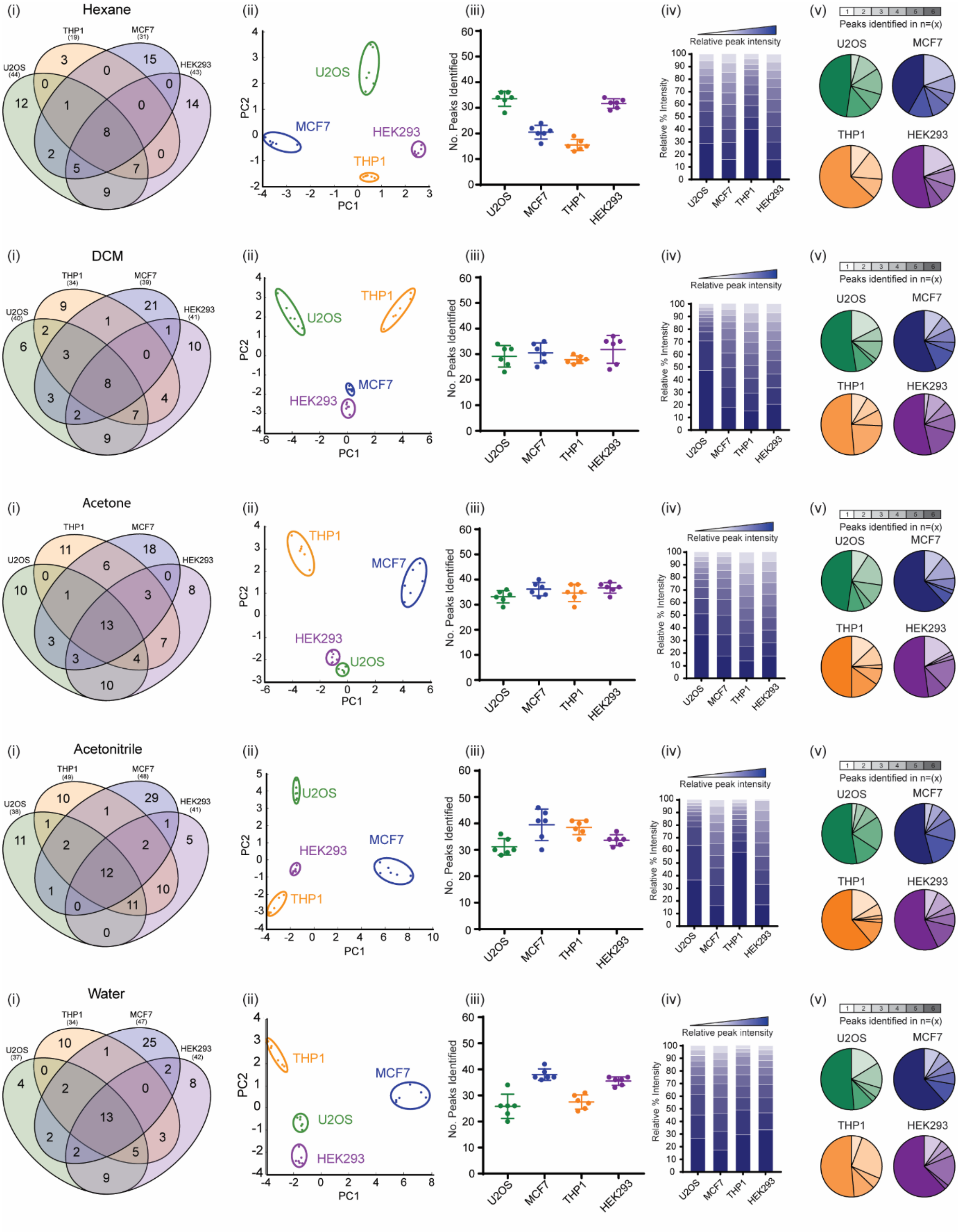

**Figure S-3: PCA plots of solvent washes with respect to cell line**. Grouping of solvent washes for each cell line indicates that each solvent interacts with the biomolecules of each cell line differently to extract unique mass profiles. Generally, apolar solvents show closer and overlapping grouping compared to PBS and water pH 7, thus indicating that apolar solvents produce more similar mass profiles.

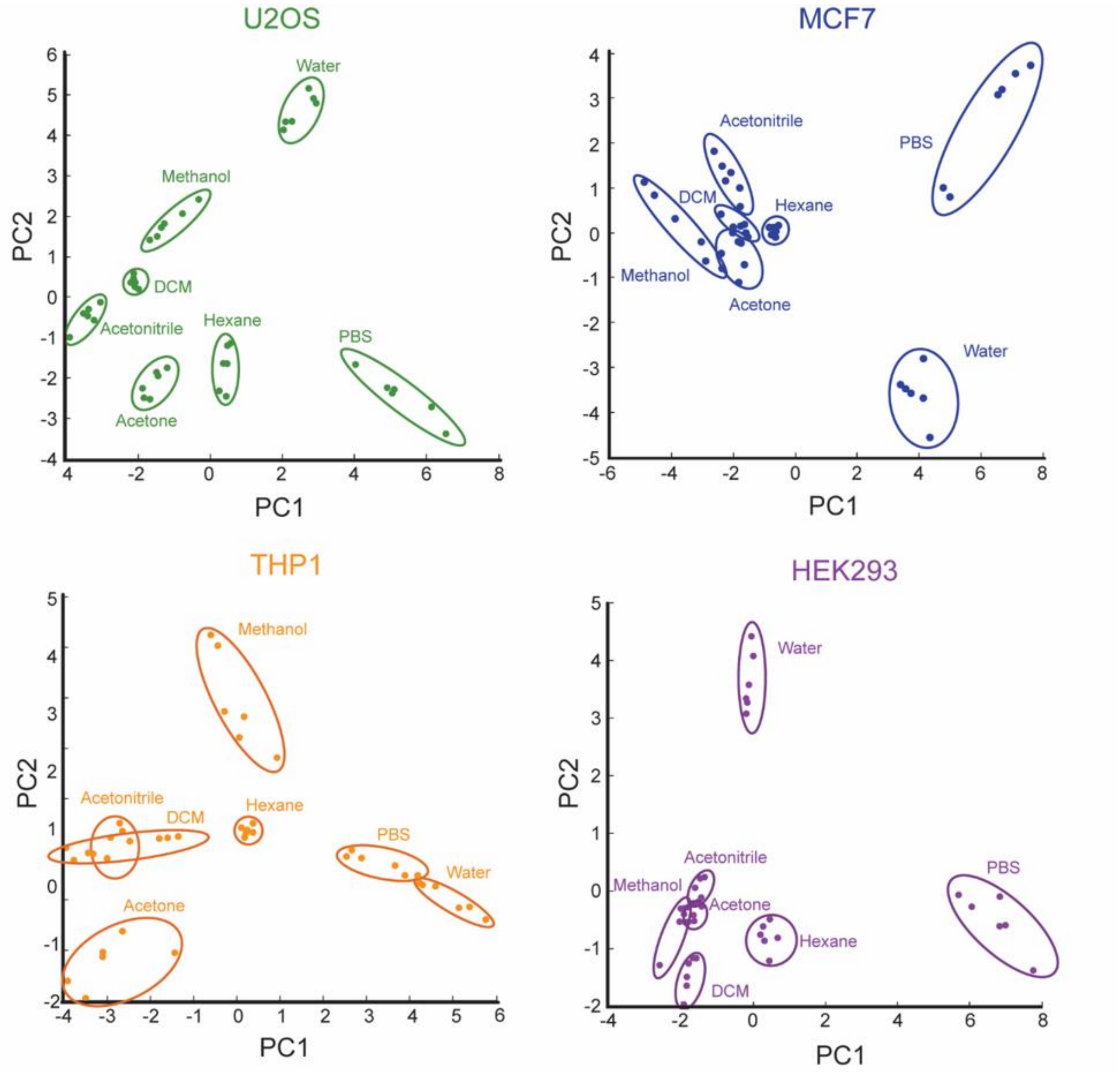

**Figure S-4: PBS wash spectra**. MALDI-TOF spectra of each of the four cell lines (U2OS, MCF7, THP1, HEK293) that were washed with PBS before subsequent analysis. For each cell line good spectra were acquired with unique profiles being generated for each.

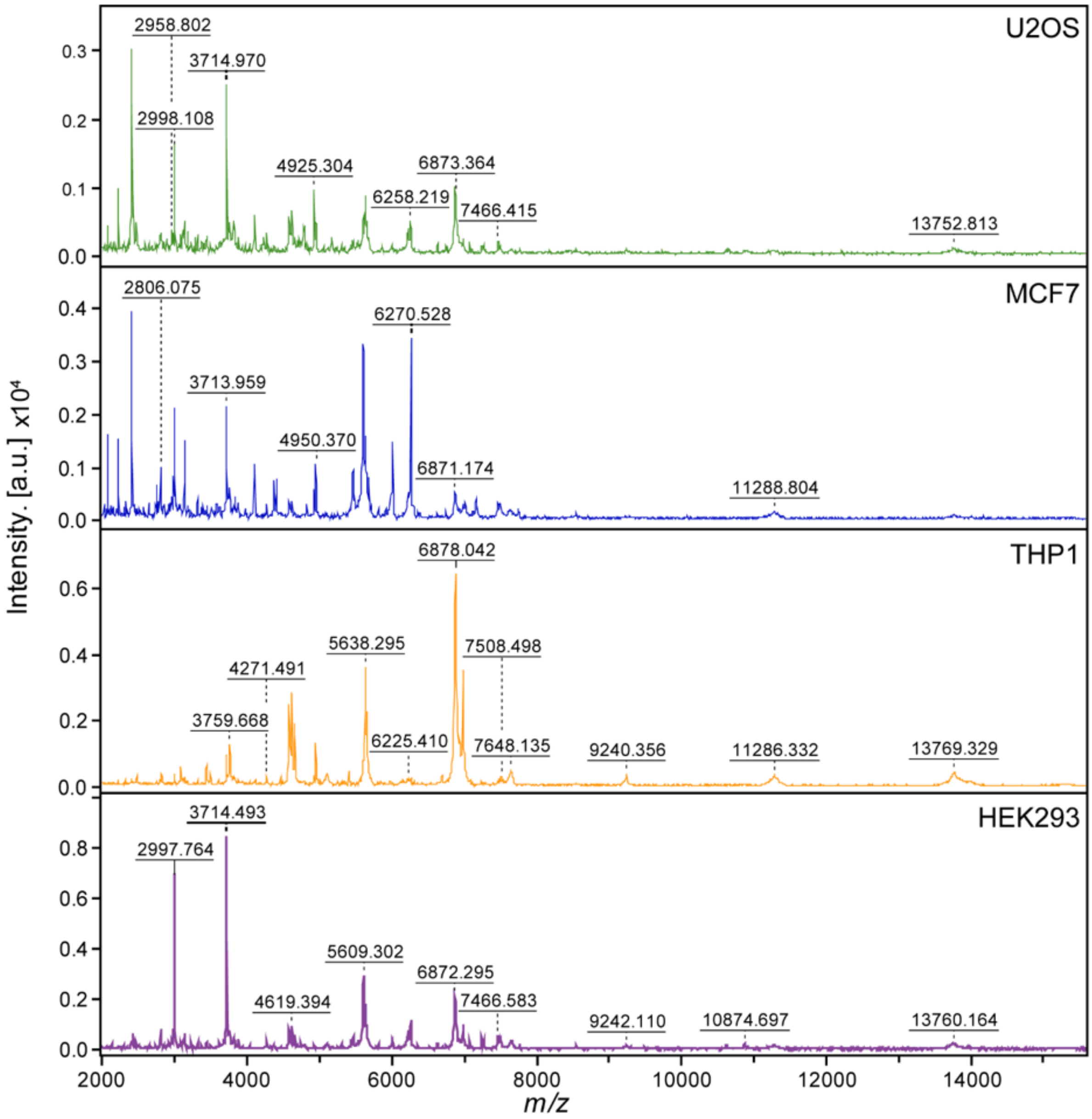

**Figure S-5: Methanol wash spectra.** MALDI-TOF spectra of each of the four cell lines (U2OS, MCF7, THP1, HEK293) that were washed with methanol before subsequent analysis. For each cell line good spectra were acquired with unique profiles being generated for each.

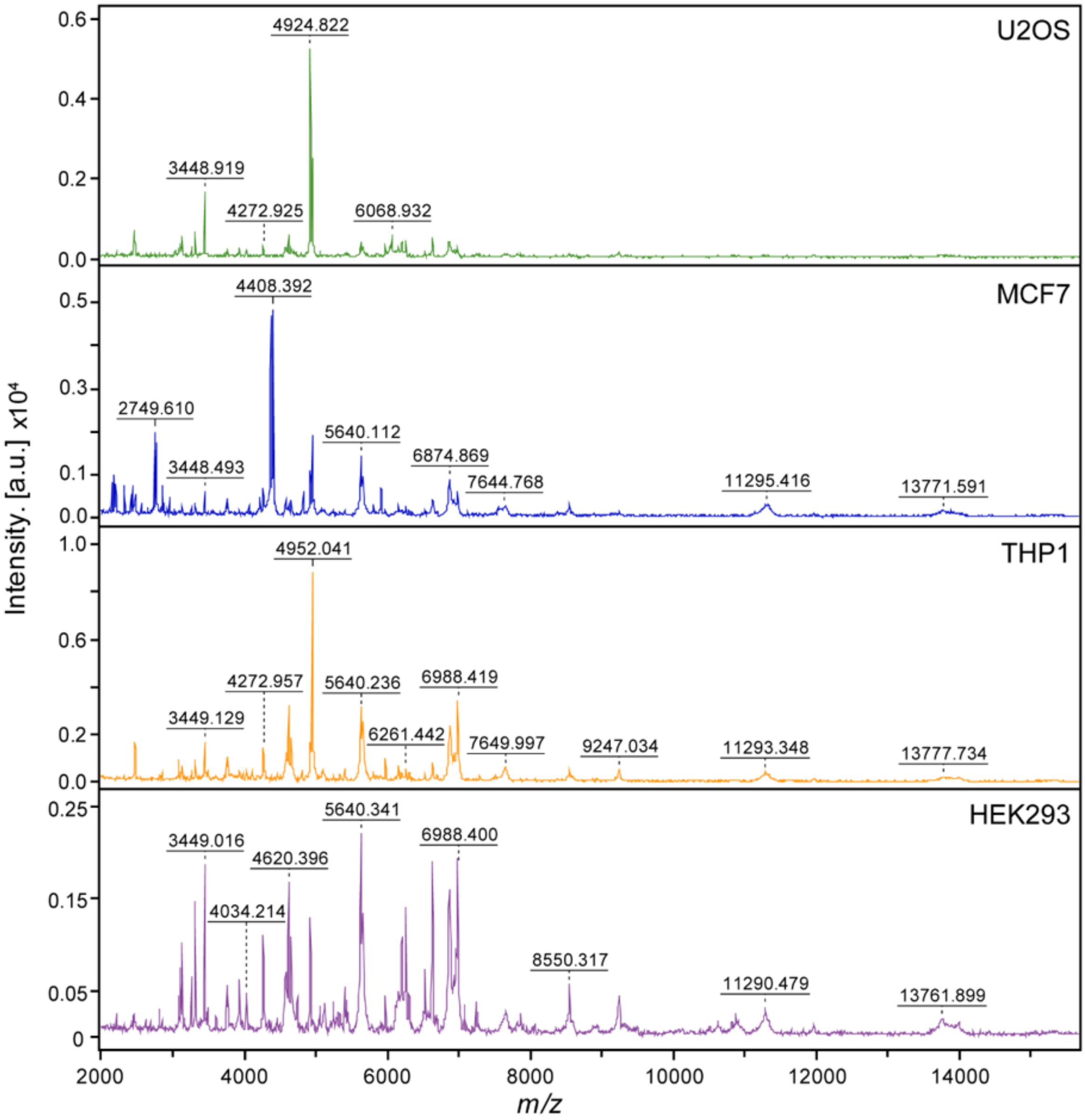

**Figure S-6 Matrix spectra.** MALDI-TOF spectra of each of the four cell lines (U2OS, MCF7, THP1, HEK293) for each of the saturated matrix conditions: α-cyano-4-hydroxycinnamic acid (CHCA), 2,5-dihydroxybenzoic acid (DHB) and sinapinic acid (SA). Unique mass spectra were acquired for each cell line when mixed with the different matrices.

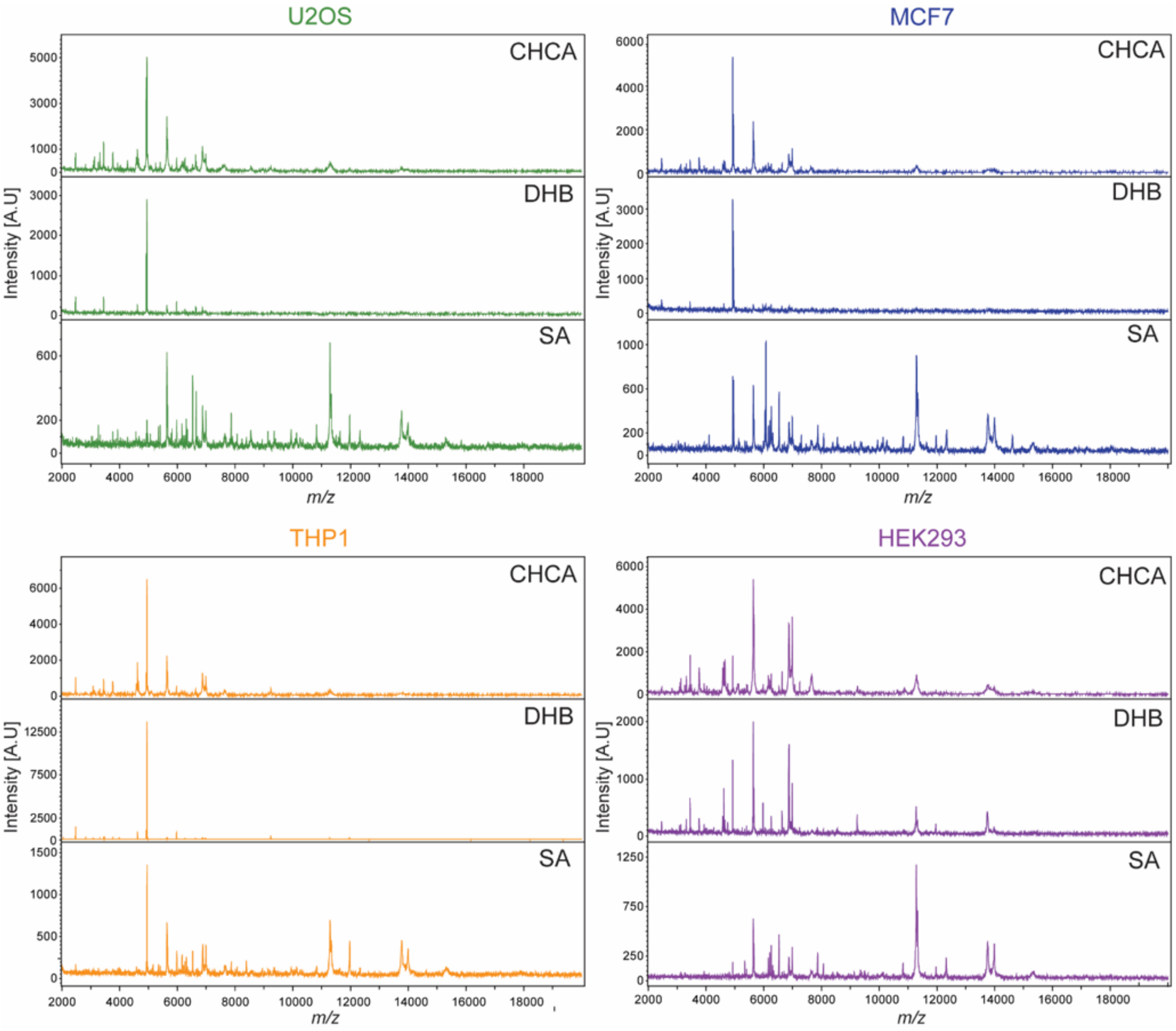

**Figure S-7 Unique features and reproducibility of each matrix condition.** Venn diagrams showing unique and common peaks identified for each of the four cell lines (U2OS, MCF7, THP1, HEK293) over six technical replicates. Bar charts displaying peak reproducibility in 50% of the technical replicates where DHB preforms significantly poorer compared to CHCA and SA.

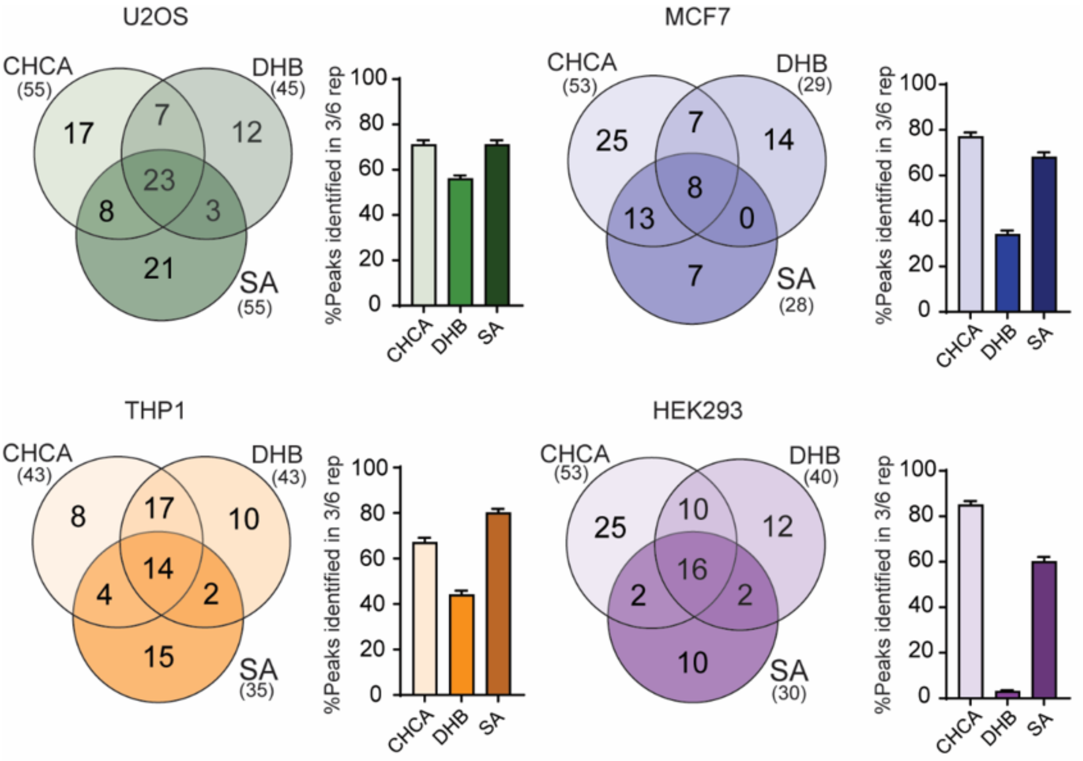

**Figure S-8 2i and 2i release spectra.** Representative MALDI-TOF spectra of each of the three individual biological replicates for 2i and 2i release cell populations.

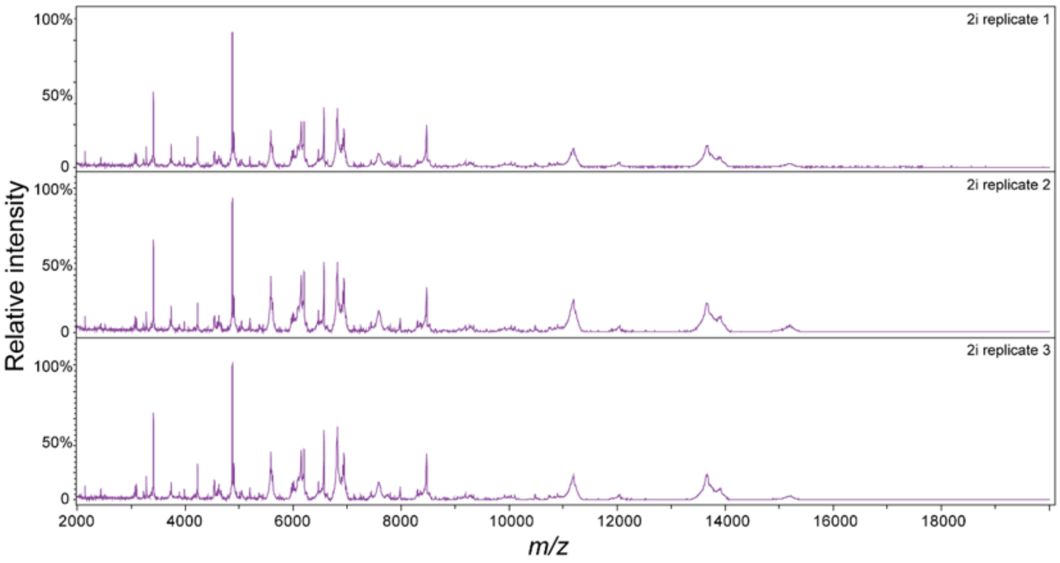


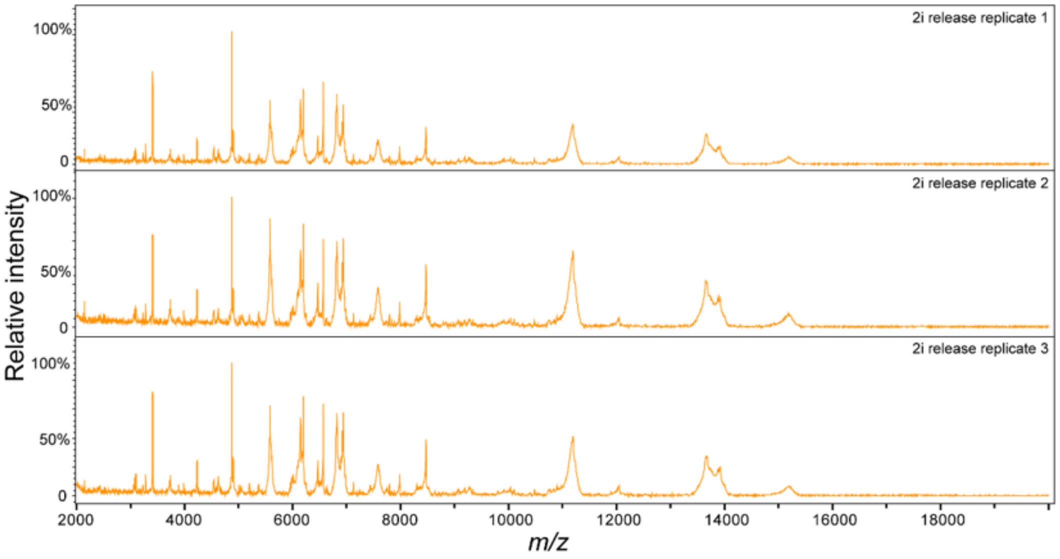
